# Nested calcium dynamics support daily cell unity and diversity in the suprachiasmatic nuclei of free-behaving mice

**DOI:** 10.1101/2021.12.14.472553

**Authors:** Lama El Cheikh Hussein, Pierre Fontanaud, Patrice Mollard, Xavier Bonnefont

## Abstract

The suprachiasmatic nuclei (SCN) of the anterior hypothalamus host the circadian pacemaker that synchronizes mammalian rhythms with the day-night cycle. SCN neurons are intrinsically rhythmic, thanks to a conserved cell-autonomous clock mechanism. In addition, circuit-level emergent properties confer a unique degree of precision and robustness to SCN neuronal rhythmicity. However, the multicellular functional organization of the SCN is not yet fully understood. Although SCN neurons are well coordinated, experimental evidences indicate that some neurons oscillate out of phase in SCN explants, and possibly to a larger extent *in vivo*. Here, we used microendoscopic Ca^2+^_i_ imaging to investigate SCN rhythmicity at a single cell resolution in free-behaving mice. We found that SCN neurons *in vivo* exhibited fast Ca^2+^_i_ spikes superimposed upon slow changes in baseline Ca^2+^_i_ levels. Both spikes and baseline followed a time-of-day modulation in many neurons, but independently from each other. Daily rhythms in basal Ca^2+^_i_ were well coordinated, while spike activity from the same neurons peaked at multiple times of the light cycle, and unveiled clock-independent interactions at the multicellular level. Hence, fast Ca^2+^_i_ spikes and slow changes in baseline Ca^2+^_i_ levels highlighted how diverse activity patterns could articulate within the temporal network unity of the SCN *in vivo*, and provided support for a multiplex neuronal code in the circadian pacemaker.

## Introduction

Circadian clocks rely on a conserved molecular mechanism expressed in virtually all body cells (1). In the suprachiasmatic nuclei (SCN) of the anterior hypothalamus, circuit-level interactions confer a unique degree of precision and robustness to circadian neuronal rhythms, which turns cell-autonomous oscillators into the central pacemaker that synchronizes mammalian physiology and behaviors with the day-night cycle (2-6). Deciphering the multicellular functioning of the SCN will thus constitute a milestone on the way from clock genes to complex, overt rhythms.

SCN neurons are coordinated but they are not always perfectly in phase. Stereotyped circadian waves in clock gene expression, intracellular calcium concentration (Ca^2+^_i_), and spontaneous firing rate of action potentials outline an approximate 6-hour phase gradient throughout SCN slices (3, 7-10). However, recent evidences challenged the rhythmic unity in the SCN, with some neurons exhibiting extreme phase lag. Phaseoids were defined as groups of neurons expressing the circadian clock protein PERIOD2 stably out of phase with their neighbors (11). Similarly, some SCN neurons appear electrophysiologically more active during nighttime, in total opposition of phase relative to the ensemble rhythm (12, 13). Last but not least, the circadian amplitude in multi-unit electrophysiological activity is dramatically downsized *in vivo* as compared to *in vitro* preparations (14), suggesting reduced neuronal synchronicity in living animals. How such a diversity in neuronal activity *in vivo* articulates within a coherent SCN network remains unclear.

Typically, fast Ca^2+^_i_ spikes driven by action potentials provide a fairly reliable estimate of neuronal activity. While circadian variations in Ca^2+^_i_ were extensively described in individual SCN neurons *ex vivo* (7, 8, 15-17), and at a neuronal population level *in vivo* (18, 19), the spatiotemporal organization of SCN Ca^2+^_i_ spikes has been barely investigated, possibly because of technical issues with BAPTA-based Ca^2+^_i_ probes (20, 21). Here we performed microendoscopic imaging of the Ca^2+^_i_ indicator GCamp6f at a single-cell resolution through a graded-index (GRIN) lens aimed at the SCN of freely behaving mice. Fast Ca^2+^_i_ spikes revealed unexpected diversity in daily neuronal activity patterns, and circuit-level signaling all around the light-dark cycle. Yet, we found that this diversity builds upon a highly coordinated wave in baseline Ca^2+^_i_ levels in the same cells. Together, Ca^2+^_i_ spikes and baseline Ca^2+^_i_ levels were independently modulated with the time of day *in vivo*, and reconciled the apparent discrepancy between functional cell diversity and unity in the SCN cell network.

## Results

After surgical preparation (see methods), mice were trained to the imaging setup for at least 3 weeks. They exhibited low levels of circulating corticosterone during their resting phase, which increased at the beginning of their active phase (Fig. S1). Moreover, longitudinal monitoring of the global GCamp6f signal from the SCN (n>10 mice) revealed daily changes, with peak and nadir in the middle of the light and dark phase (Fig. 1a,b), respectively, resembling the typical Ca^2+^_i_ rhythm recorded at the neuron population level using *in vivo* fiber photometry (18, 19). Hence, GRIN lens-implanted mice were free of chronic stress and maintained normal daily physiology under our experimental conditions.

**Fig. 1:**
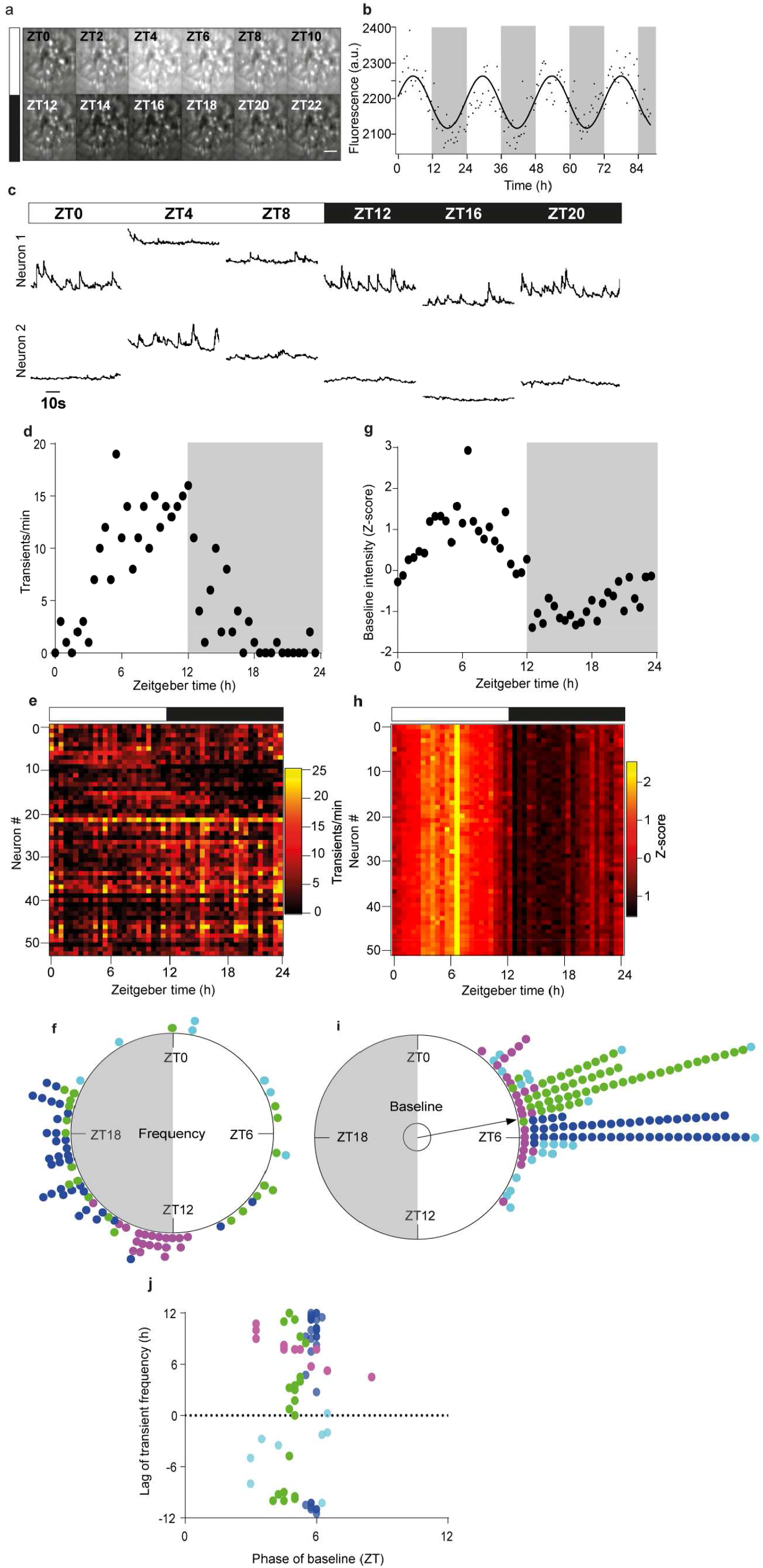
Daily diversity and unity of two Ca^2+^_i_ dynamics in individual SCN neurons *in vivo*. **a**., Time-lapse projection of GCamp6f fluorescence monitored every two hours from zeitgeber time ZT=0 to ZT=22, through a GRIN lens aimed at the SCN of a freely-moving mouse. Scale bar, 100 µm. **b**, Daily variations in global GCamp6f signal intensity from one representative SCN field of view. The solid line represents the best cosine fit of experimental data points, measured every 30 minutes over four consecutive day-night cycles. **c**, Longitudinal one-minute recordings of GCamp6f fluorescence over one day-night cycle, from two SCN neurons within the same field of view. **d-i**, Daily variation in spike frequency **(d-f)** and GCamp6f baseline **(g-i)**, in SCN neurons. Example from one representative neuron **(d, g)**, heat maps obtained from 52 neurons recorded in the same SCN field of view **(e, h)**, and Rayleigh plots of daily peak phases measured from four different mice **(f, i)**. Each color represents neurons from one mouse. The inner circle represents the statistical threshold (p<0.05) for the mean vector of the circular distribution of the aggregate data. The white and black boxes depict the light and dark phase, respectively. **j**, Phase-lag relationship between spike frequency and the GCamp6f baseline.

All the recorded SCN neurons displayed short-lived spikes in GCamp6f (Movie 1), superimposed upon slow-evolving changes in basal fluorescence (Fig. 1c). Remarkably, some neurons were more active during daytime and others during nighttime, even in the same field of view (Fig. 1c). To assess the temporal patterns in Ca^2+^_i_ activity, we estimated the frequency of GCamp6f spikes from 155 individual SCN neurons from four mice, recorded longitudinally over at least 24 hours, for 1 minute every 30 minutes. Spikes could occur at all times of day, with 38-69% of the neurons exhibiting a significant daily organization in spike frequency (Fig. 1d-f, Fig. S2-5), as assessed by non-parametric JTK-Cycle analysis (22). The large dispersion of peak activity phases from these neurons (p>0.1 Moore’s modified Rayleigh test, Fig. 1e,f) revealed unforeseen diversity of daily patterns in SCN neuronal activity, and underscored the relevance *in vivo* of night-active neuron populations previously reported in slices (12, 13).

Importantly, the daily organization in basal Ca^2+^_i_ in the same SCN neurons differed markedly from that in fast spiking activity. Up to 145 out of the 155 recorded neurons (70% to 100%) displayed a significant daily pattern in baseline GCamp6f levels that always reached a maximum during daytime (Fig. 1g-i, Fig. S2-5), in phase with the global GCamp6f signal. First, the overall phase difference between all four SCN (p<10-7, F(3, 141), Watson-Williams F-test) suggested subtle inter-individual variability, or topographical heterogeneity that was reminiscent of the phase gradient in circadian Ca^2+^_i_ levels observed in SCN slices (7, 8). However, the narrow phase distribution of maximum baseline GCamp6f levels, centered around zeitgeber time ZT=6 (with ZT=0 at lights on 95% confidence interval [ZT=03:28, ZT=07:11], p<0.001 Moore’s modified Rayleigh test), underscored the remarkable unity of daily changes in baseline Ca^2+^_i_ levels in individual SCN neurons. This resulted in a stratified phase-lag relationship with the daily variation in spike frequency (Fig. 1h), indicating that fast Ca^2+^_i_ spike activity and basal Ca^2+^_i_ levels in SCN neurons were modulated *in vivo* with the time of day, independently from each other.

To gain further insight into the contribution of spikes to the daily Ca^2+^_i_ rhythm in SCN neurons, relatively to baseline changes, we measured the amplitude of changes in GCamp6f fluorescence in each one-minute activity recording (Fig. 2a). This parameter followed a daily organization in about half (33-52%) of the recorded neurons (Fig. 2b-d, Fig. S2-5), correlated with spike frequency (Spearman r=0.94, p<0.0001) (Fig. 2e). Spiking activity was generally one-to-two orders of magnitude smaller than the variation in basal GCamp6f fluorescence measured over 24 hours in the same neuron (Fig. 2f). Therefore, spikes contributed moderately to the daily rhythmicity in fluorescence, regardless of whether they occurred at high or low baseline levels (Fig. 2g,h). These data indicated that slow changes in basal Ca^2+^_i_ levels and fast Ca^2+^_i_ spikes were nested dynamics in SCN neurons, with spikes providing cellular diversity to the coherent multicellular rhythm in baseline. Together, these Ca^2+^_i_ dynamics supported daily functional unity and diversity in the SCN cell network *in vivo*.

**Fig. 2:**
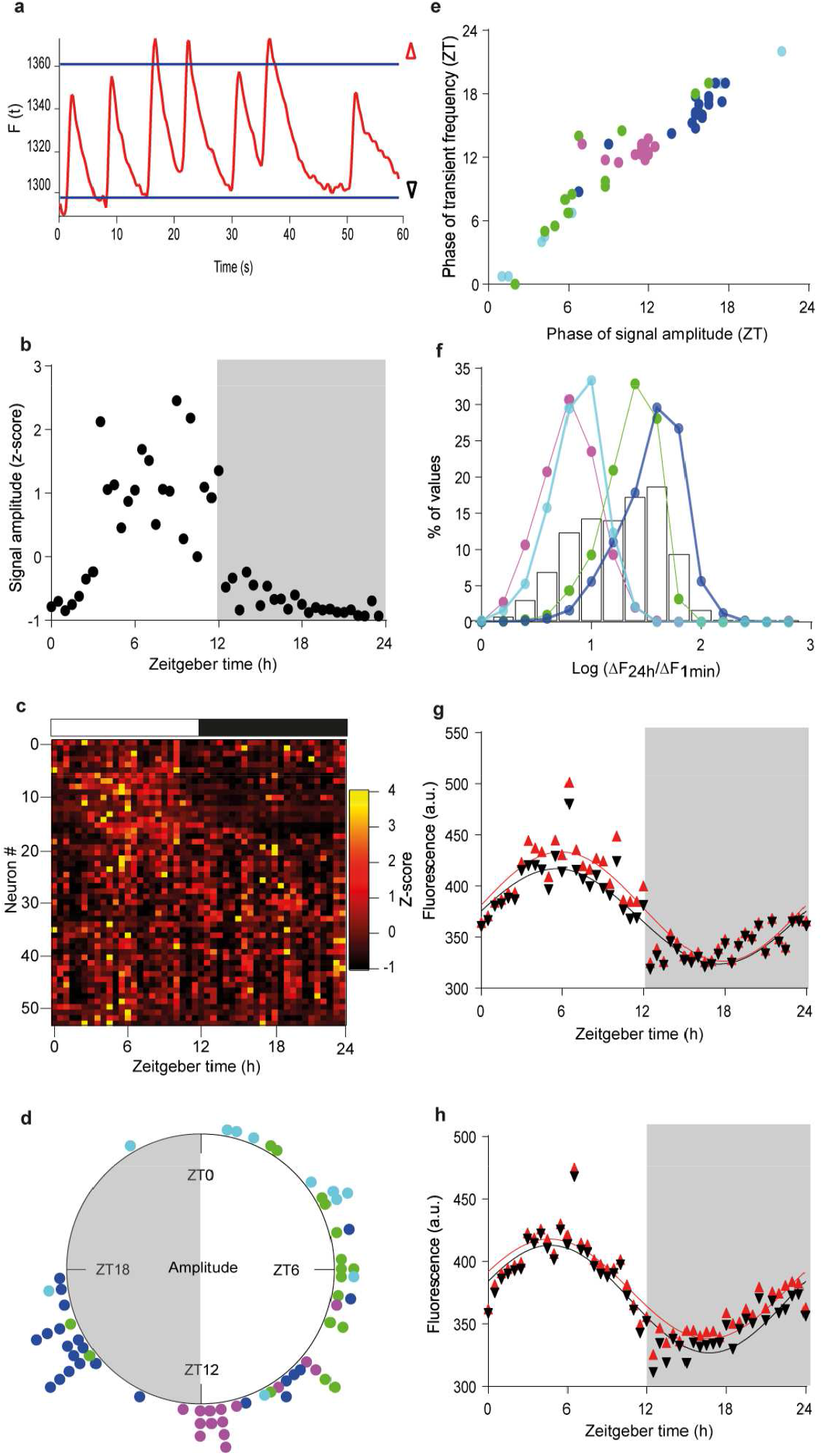
Small Ca^2+^_i_ spikes built upon large daily changes in basal Ca^2+^_i_ levels. **a**., One-minute recording of GCamp6f fluorescence, with thresholds for the 5^th^ and 95^th^ percentiles (baseline and topline, black and red triangles, respectively) used to calculate the signal amplitude. **b-d**, Daily variations in GCamp6f signal amplitude in the same neuron **(b)** and same SCN field **(c)** as in Fig. 1. **d**, Rayleigh plot of daily peak phases in signal amplitude for the aggregate data. **e**, Phase relationship between spike frequency and signal amplitude. **f**, Distribution of the ratio values calculated, for each one-minute recording, between the daily amplitude in baseline GCamp6f (ΔF_24h_) and the signal amplitude over one minute (ΔF_1min_), for each mouse SCN (colored lines) and the aggregate data (black bars). **g, h**, Daily profiles in GCamp6f baseline and topline (as depicted in a) from two representative SCN neurons, with higher signal amplitude during the day **(g)** or night **(h)**. The solid lines represent the best cosine wave for each dataset.

Next, we explored the spatiotemporal organization of GCampf spikes at the multicellular scale to question the circuit-level significance of the fast Ca^2+^_i_ activity in the SCN. We found that fast GCamp6f spikes could occur within the same time frame in clusters of SCN neurons that partially overlapped each other (Fig. 3a, Movie 1). This co-activity corresponded to noticeable changes in the global field signal (Fig. 3b), indicating that many neurons were recruited in and out of focus at the same time. The extent of co-activity clusters in the field of view, as expressed in cell.clusters/cell/min, correlated positively to the density of neuronal activity (Spearman r=0.71, p<0.0001, Fig. 3c), and spread evenly at all times of the light-dark cycle (p>0.5, Moore’s modified Rayleigh test, Fig. 3d). Since this quantitative assessment involved only co-activity detected above chance threshold (see methods), the correlation between co-activity and activity density resulted more likely from deterministic mechanisms, rather than random Ca^2+^_i_ spike coincidence in highly active neurons. These data suggested that circuit-level signaling involving Ca^2+^_i_-dependent transduction pathways synchronized SCN neurons all around the clock.

**Fig. 3:**
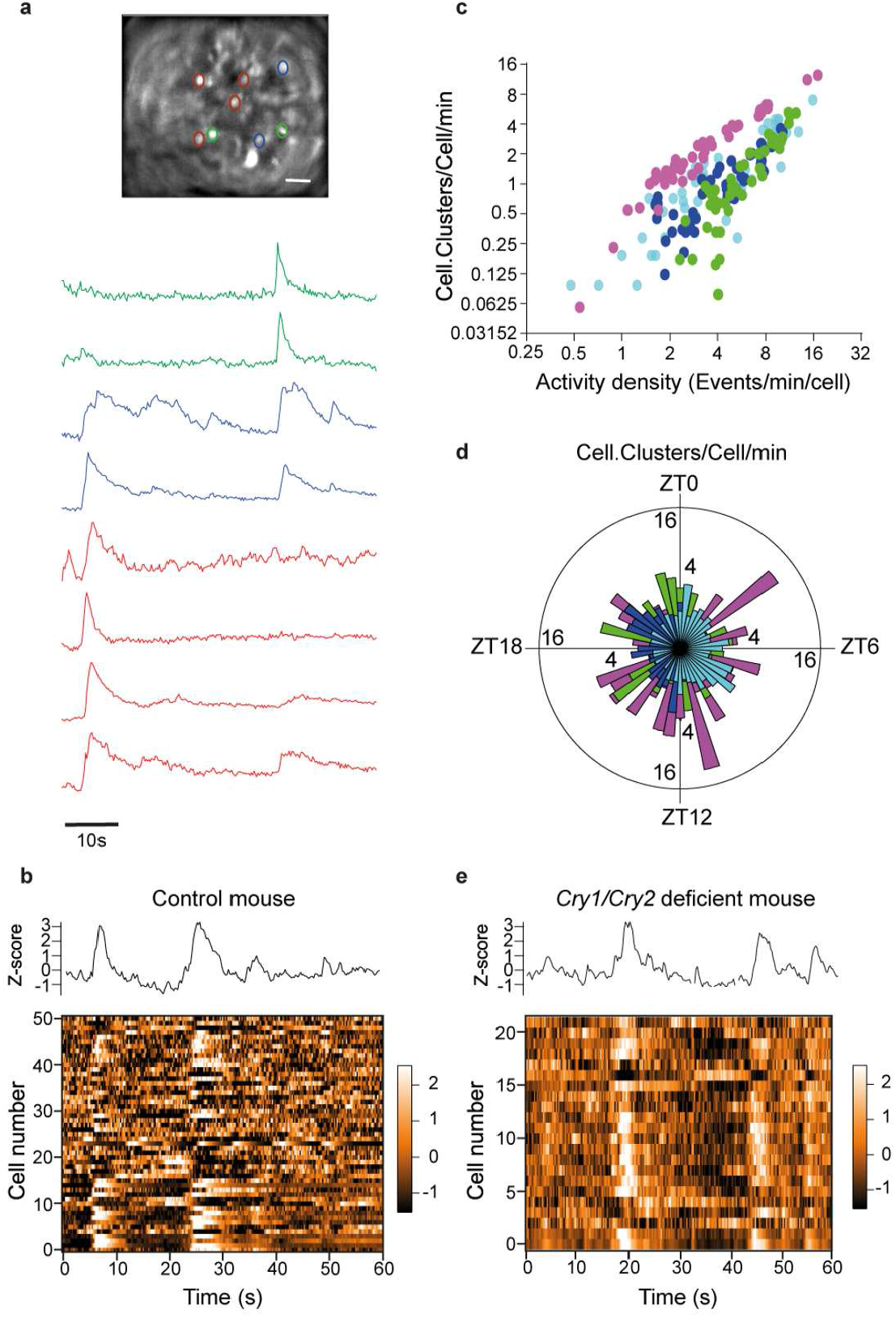
Synchronous fast Ca^2+^_i_ spikes depicted clock-independent cell-cell signaling in the SCN. **a**, One-minute recordings of GCamp6f fluorescence from SCN neurons in a same field of view. Note the occurrence of synchronous spikes in neuron subsets (delineated by three different colors). Scale bar, 100 µm. **b**, Global GCamp6f fluorescence (upper panel) and heat map of signal intensity from individual neurons (lower panel), recorded from a control mouse SCN, during one minute. **c**,**d**, Extent of clusters of co-activity as a function of the density in activity **(c)** and time of day **(d). e**, Global GCamp6f fluorescence (upper panel) and heat map of signal intensity from individual neurons (lower panel), recorded from a *Cry1-/-Cry2-/-* mouse SCN, during one minute.

To address the mechanisms underlying the concerted activation of SCN neurons, we interrogated the multicellular organization of Ca^2+^_i_ spikes under the conditions of an altered circadian clock. First, clusters of co-activity persisted during jetlag, when the global GCamp6f rhythm was progressively resetting to a new light phase (Fig. S6). Second, we conducted an independent set of GCamp6f measurements in the SCN of globally clockless *Cry1-/-Cry2-/-* mutant mice (n=4 mice). As in control mice, we observed large groups of co-active neurons that produced global events in the whole field signal (Fig. 3e). Hence, circadian clock-independent Ca^2+^_i_ signaling promoted extensive cell-cell coordination in the SCN of free-behaving mice.

## Discussion

Individuating part-whole relations in neural circuits has become an important challenge (23, 24). Our results uncovered an unprecedented relationship between fast and slow Ca^2+^_i_ dynamics that supported diversity in neuronal activity within a temporal network unity, in the SCN of free-behaving mice (Fig. 4).

**Fig. 4:**
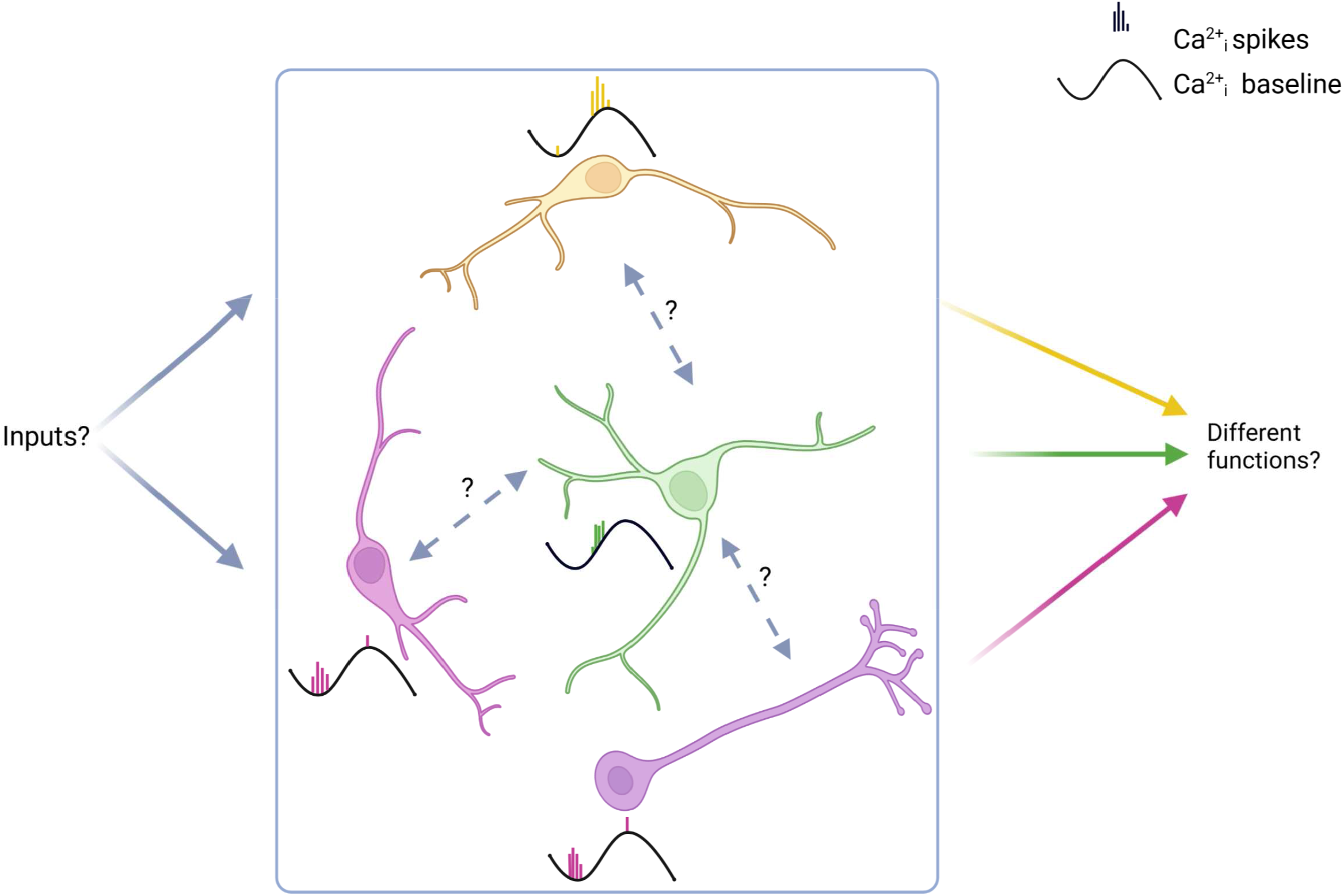
Schematic summary of the unity and diversity of Ca^2+^_i_ dynamics in the mouse SCN *in vivo*. Highly coordinated daily waves in baseline Ca^2+^_i_ levels (black sine lines) bear multiple patterns in fast Ca^2+^_i_ spikes (colored histograms), revealing neurons preferentially active during daytime (yellow and green) or nighttime (purple). Fast Ca^2+^_i_ spikes also revealed extensive clock-independent intercellular signaling, involving circuit-level interactions between SCN cells and/or coordinated inputs from other brain regions. The combination of fast and slow Ca^2+^_i_ dynamics provides support for a multiplex neuronal code to pace different body rhythms. Created with BioRender.com

At the single cell level, fast Ca^2+^_i_ spikes underscored the wide phase distribution in daily activity profiles, with a surprisingly high proportion of night-active neurons in our conditions, which somehow corroborated the elevated action potential firing rate from the SCN of live animals during nighttime (14). At the multicellular level, fast Ca^2+^_i_ spikes provided evidence of synchronization that might contribute to the unity of daily rhythms in baseline Ca^2+^_i_. Synchronized Ca^2+^_i_ spikes appeared at all times of day, and persisted under circadian clock disruption, in support of clock-independent circuit-level properties of the circadian pacemaker (10, 25). These results highlighted the relevance of Ca^2+^_i_ signaling across multiple timescales in the SCN.

The differential modulation of Ca^2+^_i_ spikes and basal Ca^2+^_i_ with the time of day suggested their relying on different mechanisms. Fast Ca^2+^_i_ spikes in SCN neurons were typically associated with electrophysiological firing, though in a non-linear relationship as individual spikes occurring at high frequency happen to sum up in amplitude (21). The correlation observed here between Ca^2+^_i_ spike frequency and signal amplitude was consistent with this assumption. Conversely, circadian Ca^2+^_i_ rhythmicity, which we found was chiefly shaped by variations in basal Ca^2+^_i_, was reportedly dependent on intracellular stores and resistant to blockers of voltage-dependent channels (16, 17). Hence, action potential-driven fast Ca^2+^_i_ spikes of low amplitude might built upon large and slow waves in daily release from internal Ca^2+^_i_ stores, although the source of each Ca^2+^_i_ dynamics likely differs among the various SCN neuronal subtypes (8, 26, 27).

Ca^2+^_i_ is a versatile second messenger that encodes a large array of cell functions (23, 28). Here, fast Ca^2+^_i_ spikes offered a tremendous extension of the repertoire of Ca^2+^_i_ signaling in the SCN, from the typical binary alternation of up and down states, toward the possibility of an activity-dependent multiplex code (Fig. 4), deciphered by the composite tuning of downstream molecular effectors that have already been reported in SCN neurons (29, 30).

Since the SCN are composed of multiple neuronal subtypes, based on gene expression profiles or neuropeptide content (31-35), the question arises as to whether neurons exhibiting the same Ca^2+^_i_ code belong to the same subtype and/or eventually constitute functional units gating specific body rhythms (13, 36-39). In a parallel study, Stowie et al. investigated circadian Ca^2+^_i_ rhythmicity in arginine vasopressin-expressing SCN neurons *in vivo* (40). Further experiments implementing this cell type-specific approach will be necessary to monitor and manipulate Ca^2+^_i_ signaling in identified SCN neurons coupled to specific circadian outputs.

## Materials and Methods

### Animals

All mouse experiments complied with the European Directive 2010/63/UE, and were registered under the reference APAFIS#15032-2018050918181227 v2. Male C57Bl/6J mice (5-6 weeks old) were purchased from Janvier Labs (Le Genest-Saint-Isle, France). *Cry1-/-Cry2-/-* mice (41) were produced by breeding double-heterozygous males and females from our colony maintained in a C57Bl6/J background for more than 20 generations (42). All mice were housed in ventilated micro-isolator cages, under a 12 h:12 h light:dark cycle, with free access to food and water.

### Surgery

At the age of 8 weeks, mice were anesthetized with a cocktail of Ketamine (Imalgene 500, 75 mg/kg) and Xylazine (Rompun, 10 mg/kg). A skin incision exposed the skull, and small craniotomies were made dorsal to the injection site with a drill mounted on a stereotaxic apparatus. The virus solution (1000 nl, titer ≥1013 vg/ml, #100836-AAV9 from Addgene) was injected in the SCN (ML +/-0.2, AP 0, DV -5.7) at a rate of 50 nl/minute using a microinjector-controlled syringe (micro-4, World Precision Instruments) and needle (Nanofil 33G beveled needle, World Precision Instruments). The needle was kept in place for 10 minutes following injection to allow suitable diffusion, and reduce backflow during withdraw. Mice were then implanted with a graded-index (GRIN) lens (diameter 0.6 mm, length 7.44 mm, working distance 150 µm, #AB000436 from GRINTECH), aimed 50-100 µm dorsally to the virus injection site. The lens was stabilized with dental cement (Metabond) and Kwik-Silsealant (World Precision Instruments). After 3-4 weeks, mice were anesthetized for placement of the microendoscope baseplate (#1050-004638, Inscopix), and a baseplate cover (#1050-004639, Inscopix) was used to protect the lens when not in use.

After surgery, mice were housed in individual cages, and regularly habituated to the imaging setup, with a dummy microendoscope (DMS-2, Inscopix) for at least 3 weeks. To ascertain well-being and normal daily physiology under these conditions, tail-tip blood (6 µl) was collected from 4 mice at ZT7 and ZT12 to check corticosterone levels (ELISA kit From Assaypro).

### Microendoscopic data acquisition and processing

Images were acquired with a head-mounted miniaturized microscope (nVista, Inscopix) at 4 frames per second over one minute (240 ms exposure time, 10-20% LED illumination, 1.5-2.5x gain). One-minute movies were spatially filtered (high-pass filter, cutoff at 40 µm), corrected for motion artifacts (Inscopix Data Processing Software), and saved as stacks of frames in TIFF format. Regions of interest (ROI) were defined and applied to every stack (ImageJ), and numerical data were saved as text files for further processing in Matlab (MathWorks).

### Data analysis and statistics

A Butterworth low-pass filter was applied (cutoff frequency at 1.1 Hz). After detrending, the baseline level for each one-minute recording was defined as the 5th percentile value, and the signal amplitude as the difference between the 5th and 95th values. Fast spikes were identified with a two-threshold procedure as fast (threshold 1, above the 90th percentile of all derivative values from all ROIs over the course of a 24-hour period) and large events (threshold 2: peak value above 2 standard deviations of all the raw signals from the considered ROI). The onset of spikes (derivative crossing above 0) was computed as their time of occurrence.

Ca^2+^_i_ spikes occurring in a high-activity frame in a statistically significant number of neurons delineated the clusters of coactivity. The threshold corresponding to a significance level of p<0.05 was defined as the number of activated cells in a single frame that exceeded only 5% of 500 surrogate datasets obtained by randomly transposing intervals of activity within each cell (43). The extent of clusters of coactivity was computed in Cell.Clusters/Cell/min, as the sum of cells participating in any of the coactive events in a one-minute recording, normalized by the total number of cells recorded in the field.

Non-parametric JTK-Cycle analysis (22) was used to estimate the daily phase of the GCamp6f signal parameters from datasets recorded over one 24-hour period. Circular analysis and statistics were conducted using Oriana (Kovach Computing Services). Other numerical and statistical analyses were conducted with IgorPro (WaveMetrics, Inc), Prism 7 (GraphPad Software, Inc).

## Acknowledgements

This work was supported by Centre National de la Recherche Scientifique (P.M., P.F., and X.B.), Agence Nationale pour la Recherche (ANR) grant nos. ANR-15-CE14-0012 and ANR-18-CE14-0017-01, and France BioImaging ANR-10-INSB-04 “Investments for the Future” to P.M., and a fellowship from Secours Populaire Libanais to L.E.C.H. We thank Ombeline Hoa and Pauline Campos for advices with GRIN lens implantation, and the iExplore and IPAM platforms in Montpellier for technical support. We are grateful to Robert J. Lucas for reading an earlier draft of the manuscript. Alec J. Davidson is acknowledged for sharing data prior to publication.

